# Published estimates of group differences in multisensory integration are inflated

**DOI:** 10.1101/331702

**Authors:** John F. Magnotti, Michael S. Beauchamp

## Abstract

A common measure of multisensory integration is the McGurk effect, an illusion in which incongruent auditory and visual speech are integrated to produce an entirely different percept. Published studies report that participants who differ in age, gender, culture, native language, or traits related to neurological or psychiatric disorders also differ in their susceptibility to the McGurk effect. These group-level differences are used as evidence for fundamental alterations in sensory processing between populations. Using empirical data and statistical simulations tested under a range of conditions, we show that published estimates of group differences in the McGurk effect are inflated. With a sample size typical of published studies, a group difference of 10% would be reported as 31%. As a consequence of this inflation, follow-up studies often fail to replicate published reports of large between-group differences. Inaccurate estimates of effect sizes and replication failures are especially problematic in studies of clinical populations involving expensive and time-consuming interventions, such as training paradigms to improve sensory processing. Reducing effect size inflation and increasing replicability requires increasing the number of participants by an order of magnitude compared with current practice.

## Introduction

Since different sensory modalities carry distinct sources of information about the world, integrating them provides a more reliable picture of the world. The underlying computations and behavioral manifestations of this multisensory integration have received increasing attention. Researchers have developed a variety of measures to assess multisensory integration, focused on a change in perception due to the addition of a second sensory modality. For instance, in speech perception, integrating auditory information from the talker’s voice and visual information from the talker’s face enhances speech recognition accuracy(1–3). Perhaps the most common, single measure of multisensory integration is the McGurk effect, an illusion in which incongruent auditory and visual speech are integrated to produce a new percept that matches neither of the component modalities. The original report of the McGurk effect(4) has been cited thousands of times (Figure 1A) and the illusion is used as a textbook example of multisensory integration. As a behavioral assay, susceptibility to the McGurk effect has been used to argue for important differences in multisensory integration between genders(5), in atypical human development(4, 6), across the lifespan(7), in mental health disorders(8, 9), and between cultural/linguistic groups (10–12). Amidst the enthusiasm for using the McGurk effect to study such between-group differences, recent work has highlighted large within-group variability in susceptibility to the illusion using relatively homogeneous subject pools (13, 14). However, most studies of the McGurk effect test a relatively small number of participants, with only a single published study reporting a group size of greater than 100 (15). Using empirical data and simulations, we show that high within-group variability, coupled with routine use of small sample sizes, leads to a proliferation of inflated estimates of group differences in multisensory integration. Our results explain why recent studies have failed to replicate the large reported differences between cultures(15), genders(13), and children with developmental disorders(16). Studies of multisensory integration must increase sample sizes by an order of magnitude to produce accurate and reliable estimates of group differences. Continued publication of inaccurate group differences can lead to vague, unscientific theories(17) and, worse, unscientific treatments aimed at ameliorating misunderstood clinical deficits.

**Figure 1.**
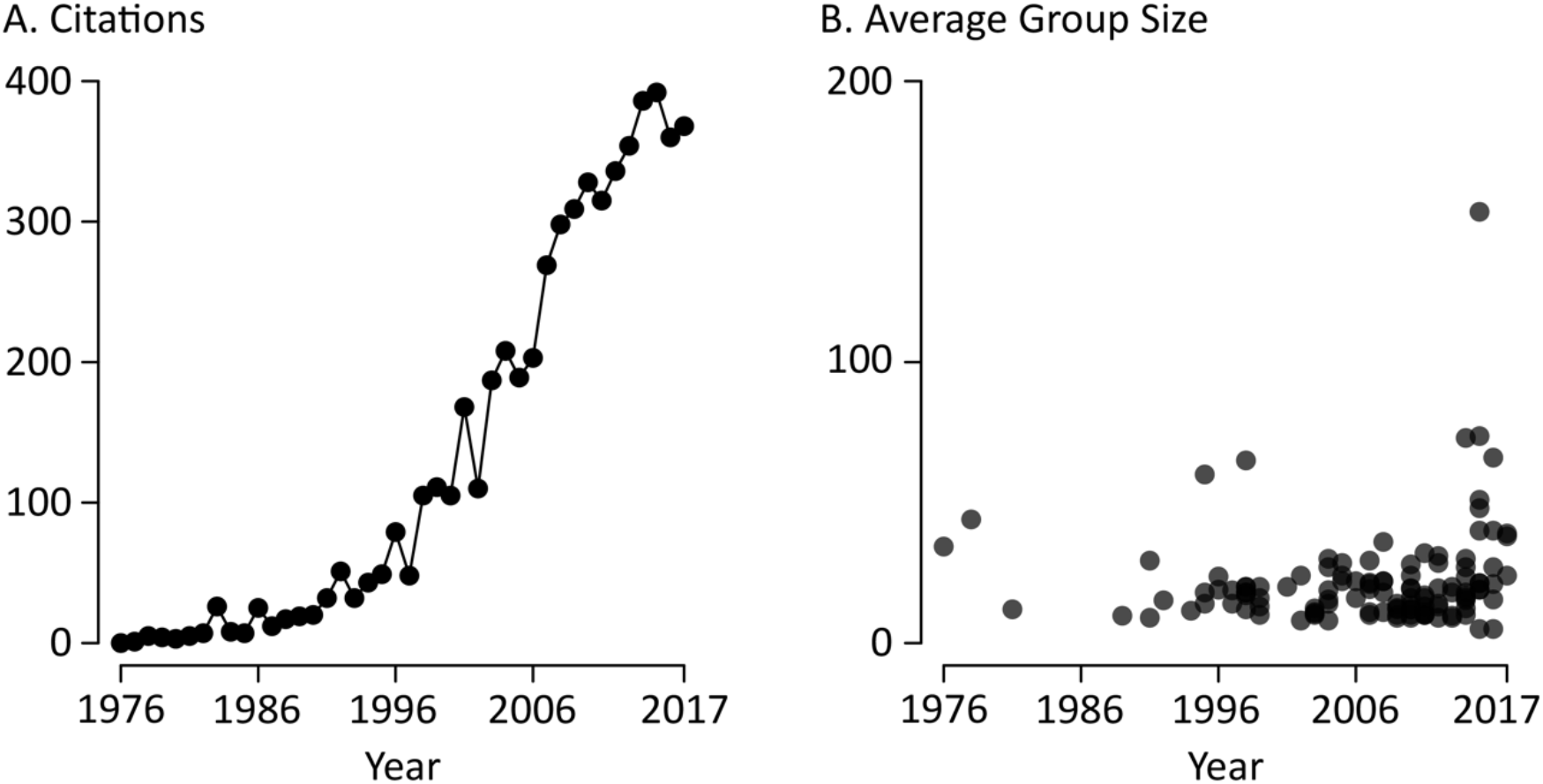
The McGurk effect in the scientific literature. **A**. The number of citations per year of the McGurk and MacDonald *Nature* paper describing the illusion, since initial publication in 1976. **B**. An analysis of the sample size (defined as the number of participants in an experimental group) in published papers on the McGurk effect.

To demonstrate how small studies can produce inflated effect estimates, we modeled the consequences of studying group differences in multisensory integration with at a variety of sample sizes. This modeling effort is made possible by a recent large-scale, in-person behavioral study, that can accurately sample the true variability within a large population of homogeneous, young healthy college undergraduates(13). For other situations, such as comparisons with clinical populations, the within-group variance is expected to be at least as large as for college undergraduates. Therefore, these simulations provide a lower bound on the sample sizes necessary to study group differences. Using empirical data rather than off-the-shelf power analysis avoids assumptions, such as population normality, that are demonstrably false for the McGurk effect(13).

### Effect inflation measured with a known true effect

To examine how sample size can influence experimental results, we used bootstrapped data from a large behavioral study and simulated population differences in McGurk susceptibility, defined as the mean percentage of fusion responses to McGurk stimuli across all possible stimuli and subjects (see *Methods* for details). In the first example, we created two populations: population A has susceptibility 45% while population B has susceptibility 55%, and thus a mean difference of 10% (Figure 2A). How likely is it for a given experiment to accurately estimate this group difference? Figure 2B shows an example experiment in which 150 subjects are sampled from each population for a total sample size of 300. With this large sample size, susceptibilities are estimated at 48% ± 3% (standard error of the mean) for population A and 60% ± 3% for population B, resulting in a difference estimate of 12% ± 9% (95% confidence interval), a reasonable approximation to the true population difference of 10%.

**Figure 2.**
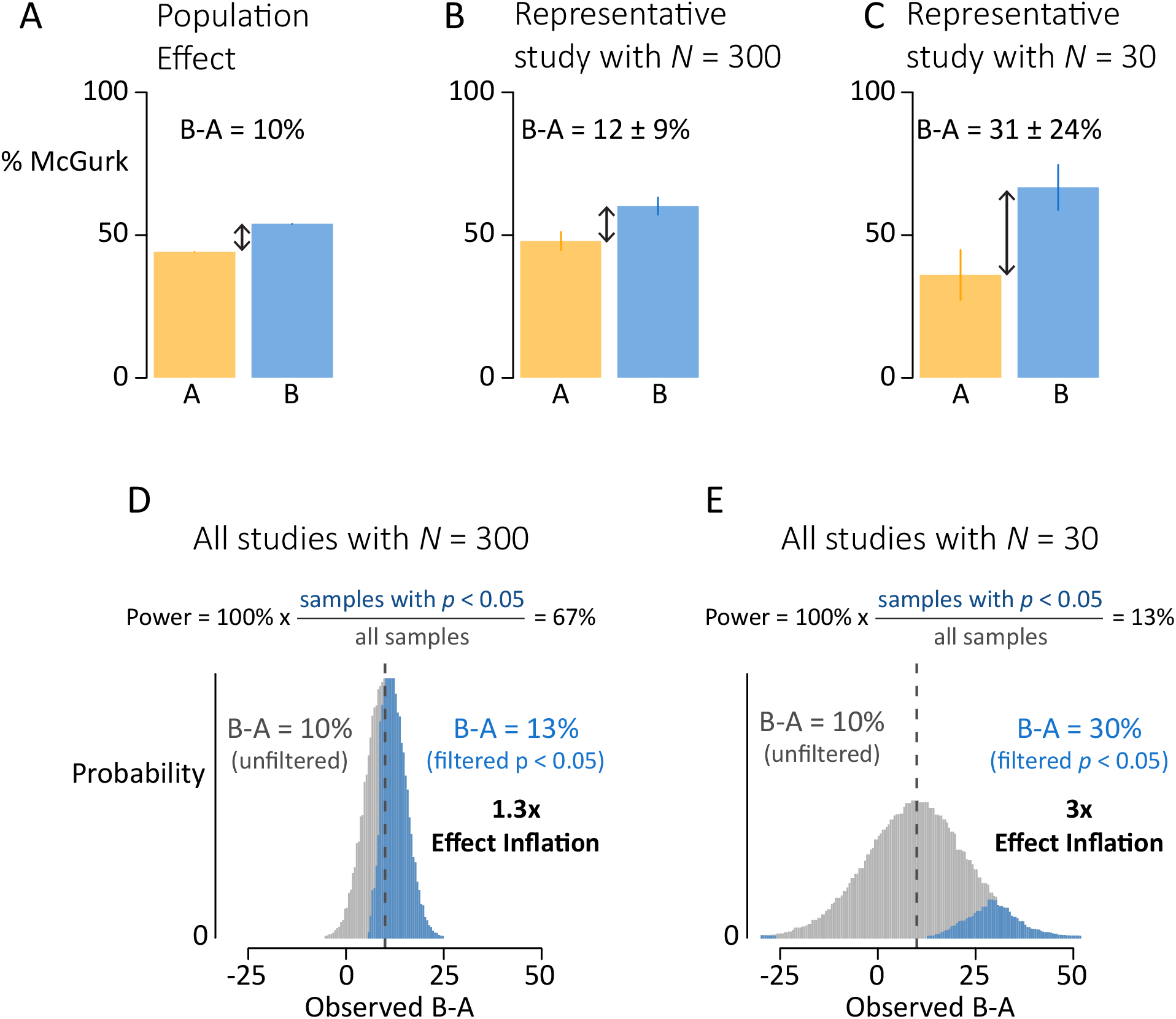
Small sample sizes lead to inflated estimates of population differences. **A**. Two example populations with McGurk susceptibility of 45% (Population A, orange) and 55% (Population B, blue), producing a true difference in McGurk susceptibility of 10%. **B**. A representative study using a sample of 300 subjects, 150 from each population. The estimated population difference is 12%, close to the true difference. **C**. A representative study using a typical sample size of 30 subjects (15 from each population). The estimated population difference is 31%, far from the true difference. **D**. Long run distribution of population difference estimates from studies with *N* = 300. Across all studies, the mean effect estimate is accurate (mean of 10%, gray bars). Considering only studies with significant results (blue bars) inflates the mean estimate to 13% because only 67% of the studies are considered (power = 67%). **E**. Long run distribution of population difference estimates from studies with *N* = 30. Considering only studies with significant results inflates the mean estimate to 30%, three times higher than the true difference on 10%.

However, a sample size of *N* = 300 is much larger than that used in published studies of multisensory integration. Figure 2C shows an example experiment with a more typical sample size of *N* = 30. This experiment results in estimated susceptibilities of 36% ± 9% for population A and 67% ± 8% for population B. The estimated population difference in this experiment is 31% ± 24%, greatly inflated from the actual difference of 10%.

This effect inflation occurs when experimenters only consider experiments that result in a significant population difference, usually defined as a between-groups *t*-test producing *p* < 0.05; this criterion can be instantiated in many ways in science, most often in the manuscript preparation and submission phase in which only significant results are included(18–20). Figure 2D shows the result of implementing this significance filter on thousands of simulated experiments with a sample size of 300. Of these experiments, 67% result in significant differences between populations and are “published”; the other 33% are discarded. The discarded studies have population difference estimates that are always *smaller* than the true difference (mean of 5%; 100% of estimates less than 10%; gray bars in Fig 1D) while the significant studies have population difference estimates that are usually *greater* than the true difference (mean of 13%; 77% of estimates greater than 10%; blue bars in Figure 2E). A weighted average of the significant and discarded experiments recovers the true population difference of 10% (13% × 0.67 + 5% × 0.32= 10%), but examining only the significant experiments biases the population difference estimate upwards by a factor of 1.3.

Figure 2E shows the results of implementing a significance filter on thousands of simulated experiments with sample size of 30. Only 13% of experiments result in significant differences between populations; the remaining 87% are discarded. The discarded studies have population estimates that are close to the true difference (mean of 7%; gray bars in Figure 2E), while the included studies have population difference estimates that are much greater than the true difference (mean of 30%; blue bars in Figure 2E). Considering only significant experiments inflates the population difference estimates by a factor of 3.

A key point is that small sample sizes not only inflate the mean effect estimate, but also increase effect estimate variance from study to study. For any particular *N* = 30 study with a true population difference of 10%, the difference estimate will vary from -16% to 36% (two standard deviations around the true difference; gray distribution in Figure 2E). This variance in effect estimation is large enough that in 3% of statistically significant studies, the population difference will be in the wrong direction: experiments will incorrectly conclude that McGurk susceptibility is higher in group A, rather than the true effect of higher susceptibility in group B. In contrast, across all *N* = 300 studies, every statistically significant study estimates the population difference in the correct direction.

### Experimental manipulations to counter effect inflation

The first example demonstrated that a true difference between populations of 10% will be vastly overestimated using procedures that are common in the literature: sample sizes of 30 and a statistical significance filter. Since true effect sizes are rarely known in advance, and are often a motivator for performing the experiment, we next examined effect size inflation at a range of true population differences. As shown in Figure 3A, effect size inflation is most extreme when population differences are small. For a true population difference of 5%, effect size inflation reaches over 7-fold for *N* = 20. In other words, on average, experiments with *N* = 20 (10 in each group) that select subjects from populations with true difference of 5% will publish effect estimates of over 35%.

In general, increasing the sample size and increasing the true population effect decrease effect inflation severity. However, for small population differences even very large sample sizes do not eliminate effect inflation: for a 5% population difference, effect inflation was more than two-fold even for sample sizes of 300. Conversely, for small *N* even a large population difference does not eliminate effect inflation. For a very large population difference of 20%, a sample size of 20 still results in two-fold effect inflation.

**Figure 3.**
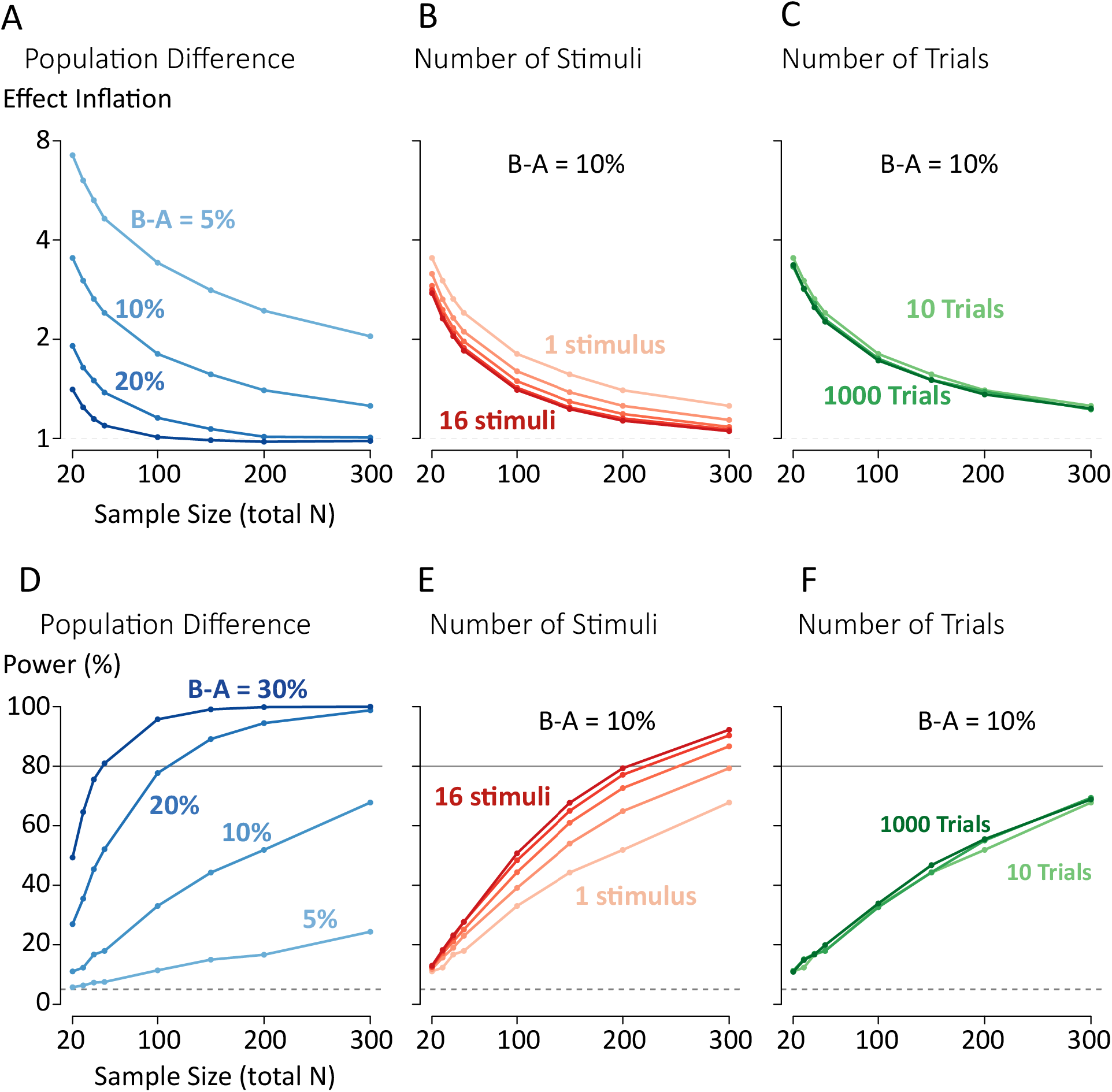
Factors contributing to effect inflation and statistical power in studies of population differences in McGurk susceptibility. **A**. Effect inflation decreases as sample size increases for a fixed population difference. Doubling the known population difference (5%, 10%, 20%, 30%; blue lines) halves effect inflation at a given sample size. **B**. Effect inflation decreases with increasing number of stimuli (1, 2, 4, 8, 16; red lines), shown for a fixed population difference of 10%. **C**. Increasing the number of trials (10, 100, 1000; green lines) has little impact on effect inflation. **D**. Statistical power (ability to detect a non-zero population difference) increases with increasing sample size and increasing population difference. **E**. Increasing the number of stimuli produces diminishing returns for increasing statistical power. **F**. Increasing the number of trials has minimal effect on statistical power.

What about other experimental manipulations? There is considerable procedural variation in the number of stimuli used to assess the McGurk effect, with some using a single stimulus, and others many more(21, 22). To examine how stimulus count influences effect inflation, we modelled the effect of increasing stimuli for a true population difference of 10% (Figure 3B). In general, increasing the number of stimuli reduces effect inflation. The effect is most pronounced when increasing the number of stimuli from 1 to 2, which reduces the effect inflation from 1.8 to 1.6 at *N* = 100. Further increases in the number of stimuli produce diminishing returns, to an effect inflation of 1.2 for 16 stimuli.

Another experimental approach is to present only a single stimulus but increase the number of trials. A typical McGurk experiment may present 10 repetitions of a single stimulus to a subject. As shown in Figure 3C, even a vast increase in the number of trials, from 10 to 1000 results in little or no decrease in effect inflation, regardless of the number of subjects tested.

### Relationship between effect inflation and statistical power

The statistical power of an experiment is defined as the probability that an experiment with a known true effect size and given sample size will result in a significant *p*-value. Power calculations implicitly incorporate a statistical significance filter, as they assume that the experimenter’s goal is to detect a difference between populations with a significance of *p* < 0.05, rather than to accurately estimate the group difference in the long run. A power of 80% is often used as a benchmark in the literature(23), as it ensures that only one in five experiments will be discarded for not being significant (assuming that the true population difference is known).

As shown in Figure 3D, large sample sizes are required for adequate statistical power across the range of population differences. Even for a very large difference between-population difference of 20% an *N* of 110 is required to achieve 80% power. For a population difference of 5% and a sample size of *N* of 300, power is only 23%.

Increasing the number of stimuli can increase power (Figure 3E), but this increase is not enough to fully overcome small sample sizes or small population differences. For a population difference of 10%, the sample size required to achieve 80% power decreases from *N* = 450 with one stimulus, to *N* = 300 with 2 stimuli, to *N* = 205 with 16 stimuli.

Changing the number of trials has minimal effect on statistical power (Figure 3F). For a single stimulus and a population difference of 10%, 450 subjects are required to achieve 80% power with 10 trials; increasing to 1000 trials only reduces the required number of subjects to 405.

### Relevance to the published literature on the McGurk Effect

Autism spectrum disorder (ASD) is frequently referred to as a disorder in which patients have impaired ability to integrate information across modalities(24). This consensus is based on behavioral studies comparing people with ASD and healthy controls. Figure 4 shows a summary of 9 of these studies, with the difference in McGurk susceptibility between ASD and healthy controls plotted against the sample size of the studies. There is wide variation across studies, ranging from 45% *less* integration in ASD (*N* = 34;(25)) to 10% *more* integration in ASD (*N* = 36;(26)). These differences across studies have been attributed to interactions between clinical diagnosis and other factors, including stimulus, gender, temporal processing abilities, or participant age. For instance, one group reported a population difference for younger children but not older children(27) while another group reported the exact opposite effect(28). We suggest that a more likely explanation than a variety of *post hoc* moderators is that small sample sizes inevitably result in high variability in population effect estimates, causing wild swings in their magnitude and sign across studies(29, 30). This contention is reinforced by noting that the ASD study with the largest sample size (*N* = 76) also reported the smallest effect, a population difference of only 3%(16). Assuming that this large study is a more accurate measure of the true effect size, we can ask what the expected effect size will be for studies with a more typical sample size of *N* = 36 (mean across the 13 comparisons made in the ASD studies). Our simulations show that applying a significance-filter to studies of this size will inflate the population difference around nine-fold, resulting in an effect estimate of 28%, similar to the observed median of 25%.

**Figure 4.**
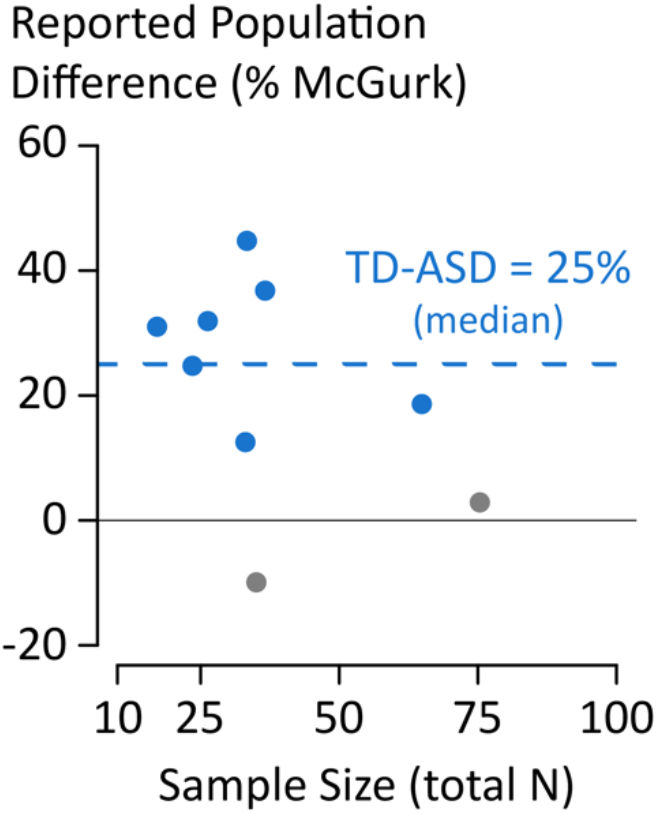
Reported population differences in McGurk susceptibility between children with autism spectrum disorder (ASD) and7 children with typical development (TD). Each symbol represents a single published? study. Statistically significant (*p* < 0.05) difference are colored in blue, non-? significant gray. Positive values indicate more McGurk susceptibility in the TD) group *vs*. the ASD group. Median across studies is 25% (blue dashed line).

A similar pattern is observed in studies of cultural differences in multisensory integration, which compare the prevalence of the McGurk effect between different linguistic or cultural populations. For instance, a number of studies have considered differences in McGurk susceptibility between speakers of tonal (e.g., Mandarin) *vs*. non-tonal (*e.g*., English) languages. Using small samples sizes, effect estimates have varied from +36% (*more* McGurk for non-tonal speakers; *N* = 24; (31)) to +17% (N = 48; (11)) to −8% (*less* McGurk for non-tonal speakers; *N* = 40; (32)). In contrast, a study with a large sample size found the smallest estimated difference: −4% (less McGurk for non-tonal speakers; *N* = 376; (15)).

Although these examples are taken from studies of population-level differences, the same problems arise in studies in which otherwise identical subjects receive different experimental manipulations, such as viewing different stimuli. In the studies of the McGurk effect we surveyed (Figure 1B), the median group size was 18 subjects.

### Relevance to other studies of multisensory integration

Problematically small sample sizes in studies of multisensory integration are not restricted to examinations of the McGurk effect. Another common assay of multisensory integration is to measure a gain in performance when a unisensory cue is provided compared to a multisensory cue. For instance, one study found *reduced* multisensory gain for individuals with ASD (N = 18) compared to typically developing (TD) individuals (N = 19) when comparing auditory-only speech-in-noise perception with audiovisual speech-in-noise perception (33), while a separate lab found (34) found *increased* multisensory gain for individuals with ASD (N = 16) compared to TD individuals (N = 14) in a multisensory temporal order judgment. In line with the latter result, but at odds with the former, another study (35) suggested reported that individuals with ASD (N = 29) integrate over a larger temporal window than TD controls (N = 17).

Although there are obvious task and stimulus differences between each of these studies, the ultimate goal of these studies (and those using the McGurk effect) is to establish generalizable estimates of group differences in multisensory integration. Critically, both multisensory speech-in-noise perception and multisensory temporal judgement tasks are known to have considerable variability even in healthy populations (36, 37). For instance, in a population of 16 healthy controls, Magnotti and Beauchamp reported a range from 70 to 300 ms in sigma (a measure akin to the temporal binding window). Just as in the McGurk effect, large inter-individual variability in healthy controls makes the measure of intergroup differences with small sample sizes unreliable in general, and inaccurate (inflated) when only significant results are published.

### Is this problem restricted to studies of multisensory integration?

Because a hallmark of many cognitive functions, including multisensory integration, is individual variation (38), large sample sizes are necessary to accurately measure a population difference or the effect of an experimental manipulation. Inadequate sample sizes are not unique to studies of multisensory integration: both the neuroscience(39) and psychology(40) literatures suffer from the same problem. Studies with small sample sizes over-estimate true effects, leading investigators in fruitless pursuit of the source of “large” effects despite failures to replicate(41). While replication failures are often attributed to *post hoc* moderators or to the inevitable experimental differences between studies, our results show that a proliferation of small studies cannot resolve conflicting results. Instead, highly-powered studies aimed at accurate estimation are the only way to provide rigorous, reliable and reproducible studies of human behavior.

### Tips for intergroup comparison studies of the McGurk effect

In this final section we summarize the results of our simulations as a series of tips for future inter-group comparisons of the McGurk effect.

#### Sample size

The major factor in determining the accuracy of the inter-group difference is the sample size. Assuming a true group difference of 20% studies with 100 subjects (50 in each group) have both reasonable statistical power (80%) and expected effect size inflation (1.2). For smaller expected effect sizes closer to 10%, 450 subjects are required for 80% power.

#### Stimulus number

In our simulations, increasing stimulus number had only moderate impacts on statistical power and effect size inflation. However, a complicating factor is that the stimuli themselves were highly variable in efficacy and the distribution of McGurk effect they elicited across subjects (13). The choice of stimulus largely depends on the goal of the study. To study *group* differences in McGurk effect, pick a small number of relatively weak (or strong) stimulus with low variation in a control population. In contrast, to study *individual* differences, it is better to use a larger number of stimuli that show high variation.

#### Trial number

In our simulations, increasing the number of trials from 10 to 1000 had no effect on statistical power. The reason for this seemingly counterintuitive result is because of the higher variation *across* subjects than *within* subjects. Increasing the number of trials an individual is given will decrease the variability in our estimate of and individuals mean McGurk perception, but it has a lesser effect on our estimate of the group-level mean. An interesting extrapolation is how *few* trials could be used and still estimate group-level differences (although individual differences would not be accurately estimated). Using only 2 trials per participant (every participant will have 0%, 50%, or 100% McGurk), a sizeable population difference of 20% can be detected at 80% power with 150 participants, compared to 100 participants with 10 trials per participant (Figure 3). Whether such a tradeoff (more participants, fewer trials per participant) is worthwhile will depend on the particulars of the hypothesis being assessed.

## Method

### Bootstrap procedure to estimate effect inflation and statistical power

We used a bootstrapping procedure to create hypothetical replication datasets based on a large behavioral dataset (*N* = 165) collected in-person from Rice University undergraduate students and described previously(13). The goal of the simulation was to determine how experimental design choices impacted statistical power (ability to reject the null hypothesis) and effect estimation (accuracy of mean estimates from studies that reject the null hypothesis). We conducted the simulations using R(42); source code is available on the authors’ website: http://www.openwetware.org/wiki/Beauchamp:DataSharing.

The simulations proceeded in a series of 7 steps:

1. Simulation parameters were set. N: number of participants (20, 30, 40, 50, 100, 150, 200, or 300), E: size of the population difference (5%, 10%, 20%, and 30%), S: number of stimuli (1, 2, 4, 8 or 16), *T*: the number of trials for each stimulus (10, 100, or 1000).
2. McGurk perception rates, *pM*, for *N* participants at *S* stimuli were sampled with replacement from the empirical data and stored in a new dataset, *D*. This procedure ensures realistic within-subject correlations across stimuli.
3. Participants in *D* were randomly assigned to group A or B, ensuring equal group sizes (group size = *N* / 2).
4. To produce the population effect, the values of *pM* was shifted by *E* for all participants assigned to group B. Because of ceiling effects, the actual size of the shift needed to be greater than the desired mean population difference. We determined these values via simulation using sample sizes of 35,000 and stimulus count of 200. Actual shift values: 9.45%, 17.25%, 33%, and 55% produce population differences of 5%, 10%, 20%, and 30%, respectively. To ensure the simulation remained stochastic, *pM* was truncated to [5%, 95%] for all participants in *D*.
5. For each participant in *D*, we calculated an *observed* McGurk perception rate *pF* for each stimulus in *S* by sampling from a binomial distribution with *T* trials and true proportion *pM*.
6. A hypothesis test was conducted to compare the *pF* between groups A and B. For single-stimulus experiments, we used a two-sample, equal-variance *t*-test. For multi-stimulus experiment, we first averaged across stimuli within each subject, and then used a *t*-test. We used a t-test rather than a linear mixed-effects model here for computational efficiency. Because of how the population effect was created (a fixed shift for all participants and all stimuli), results for the LME and *t*-test were similar.
7. To obtain long-run behavior, we repeated these steps 35000 times for each parameter combination.

To summarize the results of the simulations we used two summary measures: Statistical Power: the proportion of simulations that rejected the null hypothesis, and Effect Inflation: the mean ratio of the estimated effect magnitude (absolute value of the mean difference between groups) and the true effect magnitude, for the subset of the simulations that rejected the null hypothesis, also called the expected Type M error or exaggeration ratio(29, 30). Effect inflation provides a quantitative measure of the impact of the statistical significance filter on population effect estimates.

### Summary of population differences in children with autism spectrum disorder

To assess the impact of small sample sizes in an important area of multisensory integration research, we reviewed studies that compared McGurk susceptibility between children with autism spectrum disorder (ASD) and children with typical development (TD). We used Google scholar in the Fall of 2017 to find experimental studies comparing McGurk perception between individuals with ASD and controls. We looked for articles that cited the initial McGurk and MacDonald paper describing the illusion, using the keywords “autism”, “ASD”, and “Asperger’s”. We included only experimental studies (rather than reviews) to avoid including a specific dataset more than once. We included all studies for which group means and sample sizes were available. For studies with multiple group-level comparisons, we present them as separate data points. Two studies reported significant interactions but not group mean differences and are not included. A list of the studies used and the data collected are available from the authors’ website: openwetware.org/Beauchamp:DataSharing.

## Acknowledgments

We thank Kristen Smith for help in reviewing studies of the McGurk effect.

